# Transmission dynamics of symbiotic protist communities in the termite gut: association with host adult eclosion and dispersal

**DOI:** 10.1101/2023.09.20.558612

**Authors:** Tatsuya Inagaki, Katsura Igai, Kazuki Takahashi, Yuichi Hongoh

## Abstract

The fidelity of vertical transmission is a critical factor in maintaining mutualistic associations with microorganisms. The obligate mutualism between termites and intestinal protist communities has been maintained for over 130 million years, suggesting the faithful transmission of diverse protist species across host generations. Although a severe bottleneck can occur when alates disperse with gut protists, how protist communities are maintained during this process remains largely unknown. In this study, we examined the dynamics of intestinal protist communities during adult eclosion and alate dispersal in the termite *Reticulitermes speratus*. We found that the protist community structure in last-instar nymphs differed significantly from that in workers and persisted intact during adult eclosion, in contrast to moults between workers, in which all protists disappeared from the gut. The number of protists in nymphs and alates was substantially lower than in workers, whereas the proportion of protist species exhibiting low abundance in workers was higher in nymphs and alates. Using a simulation-based approach, we demonstrate that such changes in the protist community composition of nymphs and alates improve the transmission efficiency of whole protist species. This study thus provides novel insights into how termites have maintained mutualistic relationships with diverse gut microbiota for generations.

## Introduction

The faithful transmission of microbial symbionts between host generations is a major factor in maintaining obligate mutualism. Many insects have evolved various mechanisms that ensure high fidelity of symbiont passage during vertical transmission, including symbiont invasion of oocytes before oviposition [1] or specialized structures that contain extracellular symbionts until the egg hatches [2, 3] . However, studies of symbiont transmission have generally focused on relatively simple systems comprising one or two symbiont species [4-6]. Therefore, mechanistic insights into how a multi-species microbial assemblage (e.g., the gut microbiota) is maintained and transmitted between generations remain largely unexplored despite their prevalence and demonstrated functional importance [7].

Termites are a well-characterised example of obligate mutualism, as they are known for their associations with a variety of gut microorganisms that play an essential role in termite nutrition [8]. Except for the phylogenetically most apical family Termitidae, termites have symbiotic relationships with species-specific communities of protists in the hindgut [8]. This symbiosis likely originated approximately 130–150 million years ago or earlier in the common ancestor of termites and their sister lineage, the wood-feeding roach *Cryptocercus* [9-11]. These symbiotic protists are essential for host survival due to their roles in functions such as wood digestion [12], nitrogen fixation via their endo- and/or ectosymbiotic bacteria [12, 13], and host immunity [14, 15]. In many termite species, the protist community composition in the hindgut is highly consistent across populations [16-20], and termite-protist co-speciation has been documented [21]. These observations suggest the existence of a mechanistic basis for the faithful transmission of whole protist communities across termite generations. As intestinal protists cannot survive outside the gut of host termites, alates play a crucial role in symbiont transmission through generations in termite life history. Alates emerge from last-instar nymphs in the colony through adult eclosion and remain in the nest prior to swarming flight. To facilitate wider dispersion, their weight declines before flight, and they carry a very small number of protists [22-25], thus causing a significant bottleneck in protist community transmission. Therefore, investigating the dynamics of protist communities during this period is important in order to understand how termites overcome the significant bottleneck effect that occurs every generation. Although the dynamics of protist communities during worker development are known for some termite species, no experimental evidence of these dynamics during nymph-adult eclosion and dispersal has been reported. Workers eliminate intestinal protists before each ecdysis and regain them from surrounding nestmates by anus-to-mouth feeding (i.e., proctodeal trophallaxis) [26]. In contrast, preliminary observations reported in previous studies indicated that some protists remain during nymph-adult eclosion [20, 26-29]. This suggests that protist community dynamics exhibit a specific pattern during nymph-adult eclosion. However, whether alates receive intestinal protists from workers before dispersal remains unknown.

Here, we investigated the dynamics of the protist community from nymph-adult eclosion to dispersal in the termite *Reticulitermes speratus*. Alates of this species emerge from early April to May through eclosion from last-instar nymphs [30, 31] (Fig. 1a–d). After approximately 1 week, they disperse from the natal nest via simultaneous swarming [31] (Fig. 1e). *Reticulitermes speratus* is associated with a diverse community of protists comprising 10 species of the order Oxymonadida (*Pyrsonympha grandis, P. modesta, Dinenympha rugosa, D. exilis, D. porteri* type I–IV, *D. leidyi*, and *D. parva*) and 6 species of the phylum Parabasalia (*Trichonympha agilis, Teranympha mirabilis, Holomastigotes* sp., *Trichomonoides* sp., *Hexamastix* sp., and *Microjoenia* sp.; Fig. 1f–h). Workers harbour nearly 100,000 protists in the hindgut, and the number of each protist species varies from dozens to tens of thousands [32]. Workers of most colonies harbour all of the abovementioned protist species [17] with similar composition, suggesting faithful transmission of each protist species in *R. speratus*.

**Fig. 1.**
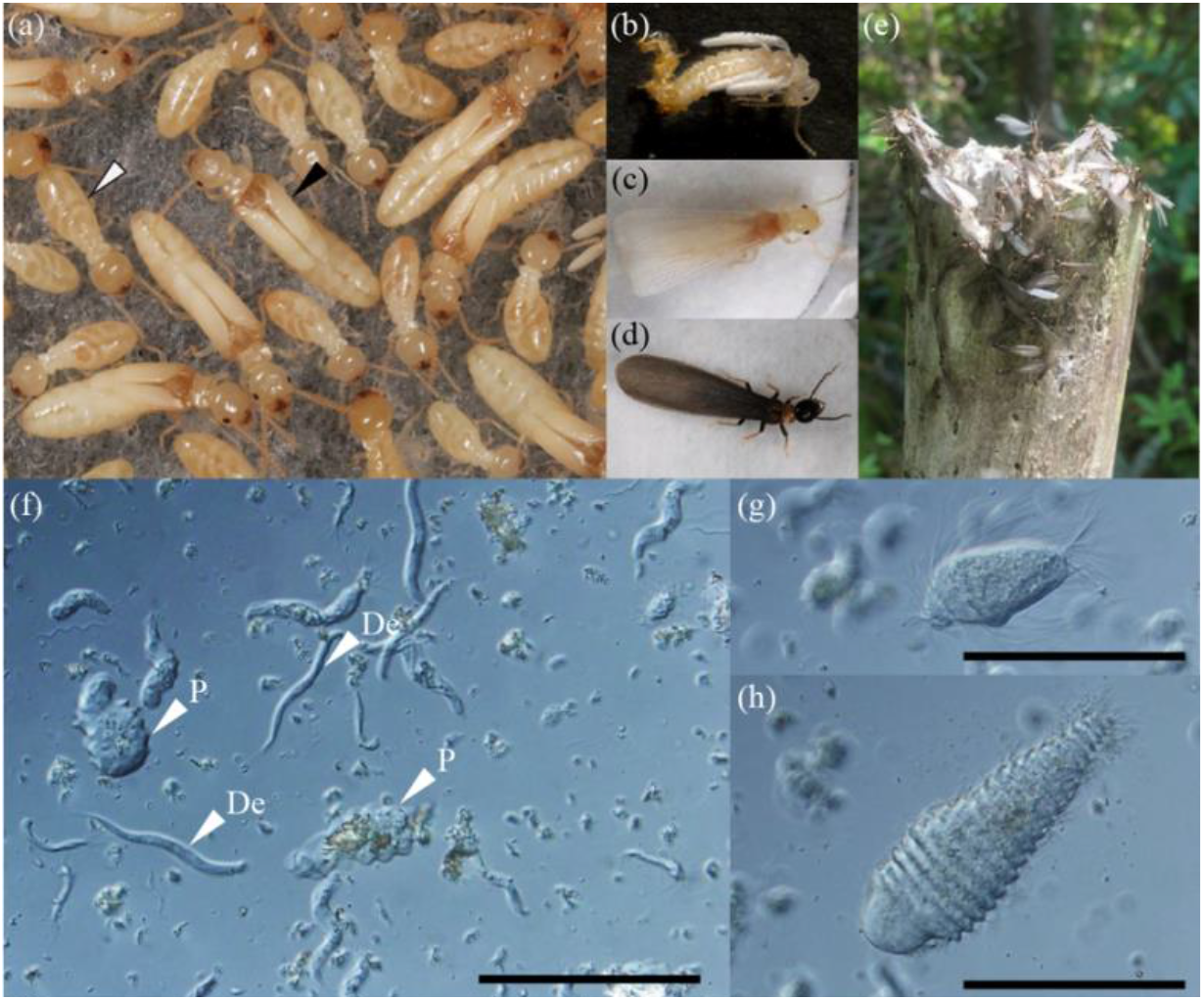
Subterranean termite, *Reticulitermes speratus*, and their symbiotic protists (a) Black and white arrowheads indicate a last-instar nymph and worker, respectively. (b-d) The process of nymph-adult eclosion. An alate (b) during, (c) immediately after, and (d) 7 days after eclosion. (e) Approximately 1 week after eclosion, alates congregate outside the nest and swarm. (f-h) Images of symbiotic protists in the gut of *R. speratus* viewed under differential interference contrast microscopy. (f) P: *Pyrsonympha* sp., De: *Dinenympha exilis*. (g) *Trichonympha agilis* and (h) *Teranympha mirabilis*. Scale bars (f-h): 100 μm.

In the present study, we first compared the changes in the protist community in the gut between the worker-to-worker and nymph-to-alate moulting stages using cell counting and 18S rRNA gene amplicon sequencing. Next, we investigated whether alates obtain protists from workers in the colony prior to dispersal. We reveal that the protist community in last-instar nymphs is a primary determinant of the community in alates because alates retain protists during eclosion and disperse without receiving any protists from workers. As the community composition of nymphs differs from that of workers, we examined whether changes in community composition contribute to transmission efficiency using a simulation-based approach. We also conducted protist species–specific fluorescence in situ hybridization (FISH) analyses and observed significant morphological changes in a protist species during adult eclosion.

## Materials and methods

### (a) Termite collection

Four colonies of *R. speratus* were collected in Japan: two in Tochigi Prefecture (designated colonies I and II) in May 2021 and Nagano Prefecture (colonies III and IV) in June 2021. Workers and last-instar nymphs were sampled from the nest wood of colonies I and II and used to investigate the dynamics of the protist community during worker-worker moulting and adult eclosion and dispersal (refer to section *(b)* below). Colonies III and IV were used to examine whether alates obtain protists from workers in the nest (refer to section *(e)* below). Last-instar nymphs were morphologically distinguished from workers and other instar nymphs by the shape of the wing pads growing from the dorsal posterolateral angles of the meso- and metathorax [30]. Total body weight, body weight without the gut, and gut weight were measured for workers, last-instar nymphs, and alates prepared in the following procedures.

### (b) Sample preparation

#### (b-1) Preparation of post-moult workers

Immediately before ecdysis, workers were sampled from two colonies. Pre-moult workers were distinguished from others by the yellowish-white colouration of the abdomen due to the gut purge [33]. The workers were isolated in 24-well plates (Stem, Tokyo, Japan) at 25°C and observed daily for moulting. Workers were collected at 2 days after moulting (moulted workers), and the absence of gut protists was confirmed by microscopic observation, as follows.

#### (b-2) Preparation of adult eclosion samples

Workers and last-instar nymphs from colonies I and II were subjected to one of two treatments: (1) isolation, or (2) being kept with workers (Fig. 2a). The sex of nymphs and alates was determined based on the morphology of the seventh and eighth sternites according to previous reports [34, 35]. In treatment (1), nymphs and workers were kept together at 20°C in several plastic cases (221 × 141 × 37 mm^3^, Mizuho-kasei, Nagoya, Japan). The cases contained brown rotten wood powder mix (brown rotten wood powder:cellulose = 1:1, [36]) and slices of Douglas fir lumber. We observed the cases daily to assess differentiation from nymphs to alates. After the first alate emerged, we extracted all of the remaining nymphs and isolated them in 24-well plates. Filter paper (Whatman No. 2, 15-mm radius, Cytiva, Marlborough, MA, USA) and 50 μL of double-distilled water were added to each well. The last-instar nymphs in the wells were observed daily, and the day of alate differentiation was recorded. Alates were extracted on the day of differentiation (A0) and 2 days (A2) after differentiation. Remaining alates were placed in the wells containing brown rotten wood powder mix and then extracted 7 days after differentiation (A7).

**Fig. 2.**
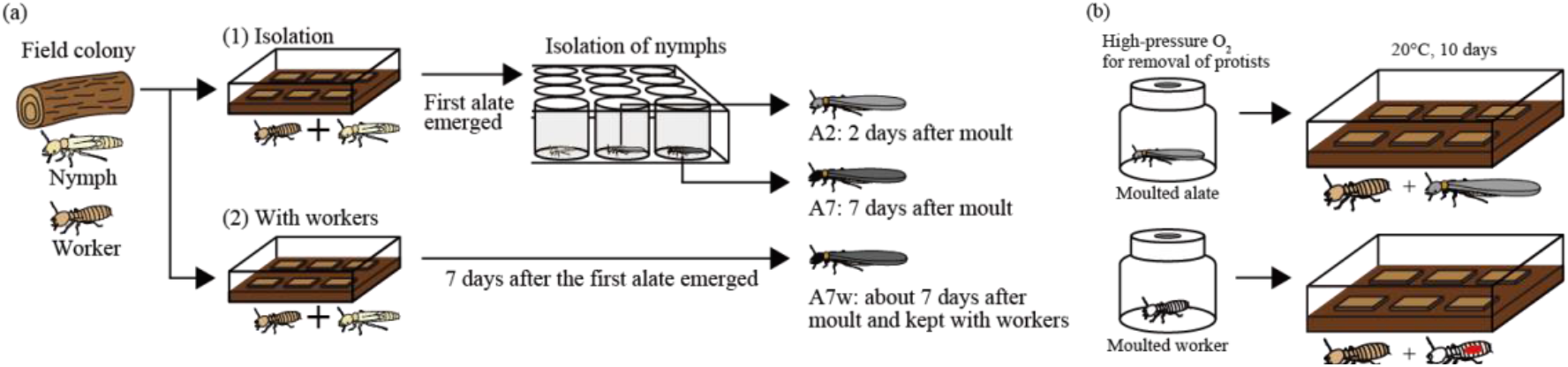
Schematic illustration of experimental procedures (a) Experimental set-up for examining the dynamics of protist communities during nymph-adult eclosion and alate dispersal. Workers and nymphs collected from two colonies were kept in (1) isolation or (2) with-worker groups. In treatment (1), nymphs were isolated after the first alate emerged in the plastic box and observed daily. Two days (A2) and 7 days (A7) after adult eclosion, alates were collected. In treatment (2), alates were collected 7 days after the first alate emerged in the box (A7w). After collection, individuals were dissected, and the protists were counted under a microscope. (b) Set-up for measuring the number of protists obtained from workers after moulting. Alates that moulted within 2 days were treated with high-pressure O_2_ (100% O_2_, 0.3 MPa for 1 h and 0 MPa for 23 h) to eliminate all protists. The alates were then kept with workers in a plastic box for 10 days at 20°C. Workers were used as a control. Moulted workers were marked with red ink to distinguish them from surrounding workers.

In treatment (2), a total of 20 male and 20 female nymphs were kept at 20°C with 500 workers in a plastic case (100 × 100 × 29 mm^3^), and nymphs were observed daily from the bottom of the case. One week after we observed the first alate differentiation, the alates were extracted as A7w. At that time, most of the alates had moved to the top of the case and were about to swarm.

### (c) Cell counting and observation of protists

The hindgut was extracted from each termite by gently pulling the tip of the abdomen using forceps. The gut contents were homogenized in Trager’s solution U [37]. The volume of solution was adjusted for each caste/group (worker: 150 μL, last-instar nymph, A2, A7, A7w: 30 μL). For all castes/groups, 20 μL of the resulting solution was loaded into a C-chip haemocytometer (NanoEntek, Seoul, Korea), and the protist cells were counted for each oxymonad and parabasalid species under a differential interference contrast microscope (IX81; Olympus, Tokyo, Japan). *Reticulitermes speratus* workers harbour up to 16 morphologically distinguishable species of protists in the hindgut [38-40], and the protists were categorized into the following categories for convenience, as some protist species can be difficult to discriminate using live specimens: *Pyrsonympha* spp. (*P. grandis* and *P. modesta*), *Dinenympha exilis, D. porteri* type III and IV, *Dinenympha* spp.1 (*D. rugosa* and *D. porteri* type I and II), *Dinenympha* spp.2 (*D. leidyi* and *D. parva*), *Tr. agilis, Te. mirabilis, Holomastigotes* sp., and small protists (*Trichomonas* sp., *Hexamastix* sp., and *Microjoenia* sp.). The large protists *Tr. agilis* and *Te. mirabilis* were counted in an area comprising 16 squares (3.2 μL) of the haemocytometer. Other protists were enumerated by counting within 4 squares at the corners (0.8 μL). The population size of each protist group was estimated, and these populations were summed to determine the total number of protists in each termite. The estimates were rounded down to the nearest integer. After determining the number of protists, the weight of the remaining body parts was determined, and the weight of the gut of each individual was estimated.

### (d) 18S rRNA gene amplicon sequencing analysis of protist community

18S rRNA gene amplicon sequencing analysis of workers, nymphs, and A7 (isolated, 7-day post-eclosion alates) was carried out using oxymonad- and parabasalid-specific primer sets, respectively. Whole-gut DNA was extracted from individual samples using Nucleo-Spin Tissue XS extraction kits (Macherey-Nagel, Düren, Germany) according to the manufacturer’s instructions, with some modifications [23]. Before the lysis step, a sterile stainless-steel bead (5 mm, Qiagen, Hilden, Germany) was placed in each tube, and the sample tubes were attached to a Vortex Adapter (24 tubes, Qiagen) and homogenized using a vortex mixer (Vortex-GENIE 2, Scientific Industries, Bohemia, NY, USA) at maximum speed for 10 min. The V3–V4 regions of 18S rRNA genes were PCR-amplified using the oxymonad-specific primers [40], with some modifications (RsOx_Sc432: 5′-GCGCAAATTACCCACTGGCA-3′; RsOx_Sc789_2Y: 5′-TTCAGCYGCGARACGCCYTG-3′), and the parabasalid-specific primers [23] (Par_18S-F: 5′-GCAGCAGGCGYGAAAC-3′; Par_18S-R: 5′-CCTACTCTCGCYCTTGATCG-3′). The PCR mixture contained 1 μL of whole-gut DNA, 1× HF buffer, 0.2 mM dNTPs, 0.5 μM primer set, and 0.4 U of Phusion high-fidelity DNA polymerase (New England Biolabs, Ipswich, MA, USA). The thermal conditions were as follows: initial denaturation for 30 s at 98°C, 25 cycles of denaturation at 98°C for 10 s, annealing at 55°C for 15 s, extension at 72°C for 30 s, and final extension at 72°C for 5 min. The PCR products were purified using Agencourt AMPure XP (Beckman Coulter, Brea, CA, USA), and DNA concentration was determined using the Qubit dsDNA HS assay kit (Thermo Fisher Scientific, Waltham, MA, USA). The purified PCR products (∼0.5 ng) were subjected to PCR amplification to attach Nextera XT indices (Illumina, San Diego, CA, USA) under the following conditions: 98°C for 30 s; eight cycles of 98°C for 10 s, 55°C for 30 s, and 72°C for 45 s; and 72°C for 7 min. Sequencing was conducted using an Illumina MiSeq platform with the MiSeq Reagent kit v3 (600 cycles).

For demultiplexing of paired-end reads, we used the clsplitseq command in the Claident package (https://www.claident.org/) [41] with a minimal quality value (Phred score) of 30 (--minqual tag = 30). Subsequently, the demultiplexed paired-end reads were quality filtered and trimmed using the dada2 v1.6 program package [42] with the following parameter settings: truncLen = c(280, 250), maxEE = c(2, 5), truncQ = 2. To compare the diversity of protists among all samples under similar sequencing conditions, the quality-filtered reads of each sample were rarefied to 10,000 read pair sets. The rarefied reads were combined (i.e., 470,000 and 450,000 read pairs in total for Oxymonadida and Parabasalia, respectively), followed by denoising, merging of paired-end reads, identification of chimeras, and sorting into single-nucleotide-level amplicon sequence variants (ASVs) using dada2. ASVs with a relative abundance of <0.1% of the total reads in a given sample and those detected in only one sample were excluded based on previous control experiments [40]. ASVs were subsequently grouped into operational taxonomic units (OTUs) based on 97% nucleotide similarity using the *DECIPHER* package in R [43]. Samples that did not yield a sufficient amount of DNA at each stage of analysis were excluded. Ultimately, 3–5 replicates were used for workers and each sex of nymphs and A7.

For taxonomic assignment of oxymonad and parabasalid OTUs, OTU sequences were aligned with the corresponding region of the near full-length 18S rRNA sequences in public databases using MAFFT [44] with the “auto” option; ambiguously aligned sites were trimmed using TrimAl [45] with the “nogaps” option. Maximum-likelihood trees were constructed using IQ-TREE v1.6.12 [46] with the TIM3+F+I+G4 and TIM2e+G4 models for oxymonad and parabasalid trees, respectively, with ultrafast bootstrap analysis [47] with 1000 re-samplings.

### (e) Removal of protists using oxygen gas

Whether alates obtain protists from workers after eclosion was determined by keeping workers and alates in which the intestinal protists were eliminated by oxygenation treatment [48] (Fig. 2b). Last-instar nymphs were placed in a petri dish with moist unwoven cloth and maintained at 20°C until they become alates by eclosion. Alates that moulted within 2 days (16 males and 20 females) were transferred into a rubber-stoppered gas chromatography vial (net capacity 30 mL, SVG-30, Nichidenrika-Glass, Kobe, Japan) for treatment with oxygen to eliminate intestinal protists (Fig. 2b). The inner bottom of each glass vial was lined with moist unwoven cloth. An oxygen spray can (Nitto Kagaku, Nagoya, Japan) was connected to the vial via an injection needle, and an oxygen concentration meter (Ichinen Jikco, Nagoya, Japan) was also connected to the glass vial via another needle with the air being allowed to flow out. Oxygen was loaded into the vials until the concentration reached approximately 100%, after which the oxygen concentration meter was disconnected and a pressure gauge (APG-1, Fujiwara Sangyo, Hyogo, Japan) connected. Oxygen was loaded into the vial again until the pressure reached 0.3 MPa. The alates were kept in the presence of high-pressure oxygen for 1 h, after which the vials were depressurized by insertion of an injection needle, and the alates were kept in the vials for another 23 h. Four males and four females were used to confirm the successful elimination of protists, and 12 male and 16 female alates were transferred and kept with 500 workers in plastic cases containing brown rotten wood powder mix with slices of Douglas fir lumber at 20°C. After 10 days, surviving alates (4 males and 7 females in each colony) were removed, and the number of protists was determined as described above section *(c)*. For a control, 44 workers that moulted within 0–3 days were treated under high oxygen conditions in the same manner as the alates. After oxygen treatment, six workers per colony were examined to confirm the absence of protists in the gut. The dorsal side of the abdomen of each post-moult worker was marked with red oil-based paint (Mitsubishi pencil, Tokyo, Japan) to distinguish it from other workers in the plastic case. The painted workers were kept with 500 workers in the plastic cases. After 10 days, 22 post-moult workers were removed, and the number of protists was determined as described above.

### (f) Simulation of vertical transmission of the protist community

A theoretical investigation was conducted to examine the potential impact of changes in protist community composition in nymphs on the transmission efficiency of the entire protist community. Using computer simulations, we modelled two transmission scenarios: random transmission to alates of protists from either (i) nymphs or (ii) workers. Sampling with replacement was performed, selecting 500 to 10,000 protist cells (in increments of 500) from a hypothetical protist community in workers or nymphs, and these were then introduced into protist-free hypothetical alates. The community compositions in the hypothetical workers and nymphs were determined as the average of empirical data from colonies I and II (Table S1). Once all protists were placed in hypothetical alates, each alate was assessed for the presence of each protist species. The transmission efficiency, that is, the proportion of alates that have all protist species after 5,000 iterations, was then calculated. The R code used for these simulations is available at https://github.com/TatsuyaInagaki2/Data_alate_protist_community.

### (g) Morphological observation of protists during adult eclosion

We designed oligonucleotide probes to specifically target 18S rRNA specific to *D. exilis* and *D. leidyi* using the probe-designing function in ARB [49]. The probe sequences were 5′-ACGCACAGGGAAAACGAA-3′ for *D. exilis* and 5′-GAATGAAGCATCGGCACG-3′ for *D. leidyi*. 6-Carboxyfluorescein (6FAM) and Texas red were attached to the 5′ ends of the probes for *D. exilis* and *D. leidyi*, respectively. The gut contents of eight individual workers and four males and four female nymphs, A0, A2, and A7w from each colony were examined. Fixation and hybridization were performed as described previously [50, 51], with slight modifications. Specimens were subjected to prehybridization in buffer (0.9 M NaCl, 0.1 M Tris-HCl) at 48°C for 15 min followed by hybridization with the probes at 48°C for 3 h. Samples were observed using an epifluorescence microscope (BX51, Olympus, Tokyo, Japan). Probe specificity was confirmed using specimens from gut samples of workers. Protist images were processed using Adobe Photoshop CC. The edges of cells were trimmed using the Quick Selection Tool function, and the area (μm^2^) and circularity (4П [area/perimeter^2^]) were determined, with a circularity value of 1.0 indicating a perfect circle. We processed 25 cells of each protist species from each caste/group (worker, nymph, A2, A7, and A7w) of the two colonies.

### (h) Statistical analysis

The total number of protists, fresh body weight, body weight without gut, and gut weight in each caste/group (worker, nymph, A2, A7, and A7w) were compared using linear mixed models (LMMs) and likelihood ratio tests (LRTs). In the LMMs, caste/group was included as an explanatory variable and colony as a random factor. To elucidate the effect of nymph and alate sex, we also ran LMMs including sex, caste/group, and their interaction as explanatory variables and colony as a random factor.

To examine the effects of colony and caste/group on protist communities, we conducted PERMANOVA (permutational analysis of variance) on Bray-Curtis dissimilarity indices calculated from cell count data. Colony, caste/group, and their interaction effect were included as explanatory variables. To elucidate the effect of sex in nymphs and alates (A2, A7, and A7w), we also ran PERMANOVA including sex, caste/group, and colony as explanatory variables. The effects of colony, caste/group, and their interaction on the proportions of *Tr. agilis* and *Te. mirabilis* in cell count data were tested using a generalized linear model (GLM) with quasi-binomial error distribution, as the data were over-dispersed.

For read counts of oxymonad and parabasalid OTUs derived from amplicon sequencing, we performed non-metric multidimensional scaling (NMDS) and PERMANOVA on Euclidean distances calculated from centred log-ratio–transformed data due to the data compositionality [52, 53]. The effects of colony, caste (worker, nymph, and A7), and their interaction were tested. To elucidate the effect of nymph and A7 sex, we also ran PERMANOVA including sex, caste, and colony as explanatory variables.

To compare the area and circularity of two oxymonad protist species, we conducted pairwise Wilcoxon tests among castes/groups (worker, nymph, A0, A2, and A7w) in each colony.

LMMs were conducted using the *lme4* package [54], and PERMANOVAs were performed using the *vegan* package [55] in R.

## Results

### (a) Persistence of protist communities during adult eclosion from nymphs to alates

Workers from the field colonies harboured a number of protists in their gut, whereas none retained any protist cells immediately after moulting (workers: 79248.6 ± 4432.9, moulted workers: 0 ± 0, mean ± SE, Fig. 3a). In contrast, the number of protists did not change significantly during nymph-adult eclosion (nymphs: 8992.1 ± 933.2, A2: 8457.0 ± 503.8, Fig. 3b). Seven days after eclosion, the number of protists significantly decreased in A7 (5193.5 ± 370.1, Fig. 3b). The decrease of protists in A7w was smaller (6770.2 ± 428.5, Fig. 3b), but the number of protist cells was not significantly different from that of A7. Among the nymphs and alates examined, the sex of individuals and interaction between sexes and castes/groups had no significant effect on the total number of protists (LMM, LRT, 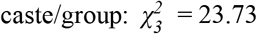, *P*<0.001; 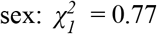, *P* = 0.38; 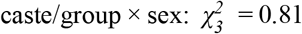, *P* = 0.85).

**Fig. 3.**
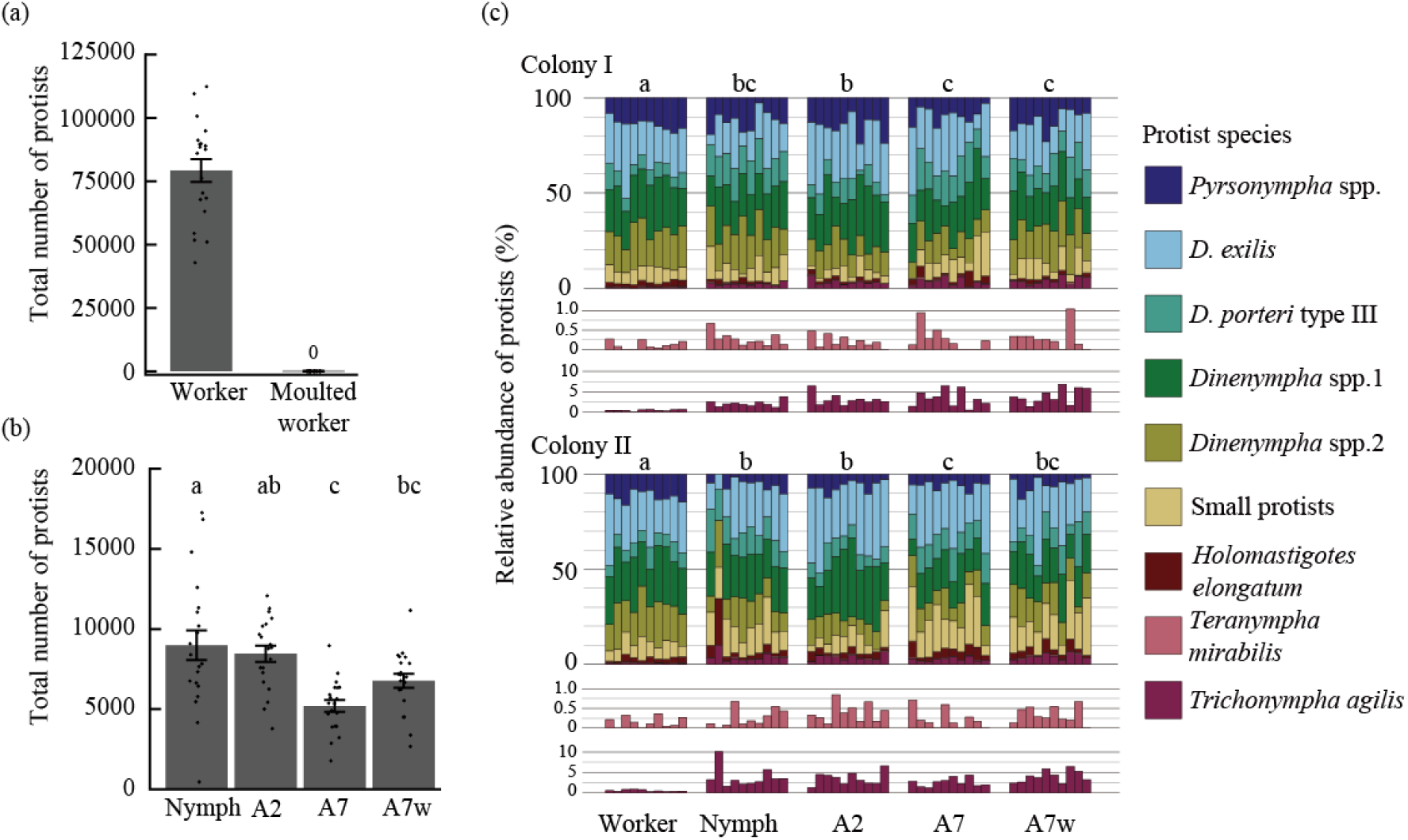
Dynamics of intestinal protist communities during alate differentiation and dispersal Changes in the number of protist cells during (a) worker moulting and (b) nymph-adult eclosion and alate dispersal are shown. Different characters denote significant differences. LMM, colony was included as a random factor, Tukey’s HSD, *P*<0.05. In (a) and (b), error bars denote the standard error, and points are individual measurements. (c) Comparison of protist communities among castes/groups. Different characters indicate significant differences between protist communities. PERMANOVAs were conducted between castes/groups in each colony (Bray-Curtis dissimilarity, 9999 permutations, Bonferroni correction, corrected *α* = 0.005). The relative abundance of *Teranympha mirabilis* and *Trichonympha agilis* in each sample is additionally shown at the bottom.

The protist community composition based on cell counts differed significantly between castes/groups and between colonies, whereas the colony and caste/group interaction effect was not significant (PERMANOVA, caste/group: *F*_*4*_ = 56.73, *P*<0.001; colony: *F*_*1*_ = 5.99, *P* = 0.004; caste/group × colony: *F*_*4*_ = 1.22, *P* = 0.274). In both colonies I and II, the protist community composition did not change before or after adult eclosion and was not affected by worker presence (Fig. 3c). PERMANOVA of nymphs and alates with colony, caste/group, and sex as explanatory variables showed no significant effect of individual sex on protist community composition (*F*_*1*_ =1.39, *P* = 0.22). When focusing on *Te. mirabilis* and *Tr. agilis*, which showed the lowest and second lowest abundance in workers, respectively, we found that proportions of both species increased in nymphs and alates (Figs. 3c, S1). The proportion increased approximately 6-fold in *Tr. agilis* (percentage proportion in workers: 0.48 ± 0.05, nymphs: 2.93 ± 0.44, A2: 3.30 ± 0.32, A7: 2.93 ± 0.36, A7w: 3.98 ± 0.37, mean ± SE, Fig. S1) and 2-fold in *Te. mirabilis* (percentage proportion in workers: 0.14 ± 0.03, nymphs: 0.27 ± 0.04, A2: 0.29 ± 0.05, A7: 0.24 ± 0.06, A7w: 0.31 ± 0.06, Fig. S1). There were no significant differences between colonies in terms of the proportions of *Tr. agilis* (GLM, LRT, 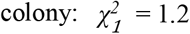, *P* = 0.27; 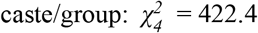, *P*<0.001; 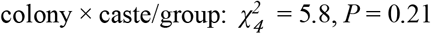) and *Te. mirabilis* (GLM, LRT, 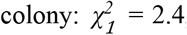, *P* = 0.122; 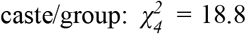, *P*<0.001; 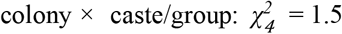, *P* = 0.83).

In the amplicon sequencing analyses of oxymonad protists, we assigned 20 OTUs to 9 species-level taxa belonging to 2 genera (*Pyrsonympha* and *Dinenympha*) (Fig. S2). For parabasalid protists, we assigned 10 OTUs to 8 species-level taxa (Fig. S3). The oxymonad community differed significantly among castes (workers, nymphs, and A7) and between colonies, whereas the interaction effect of colony and caste was not significant (PERMANOVA, caste: *F*_*2*_ =2.9, *P*<0.001; colony: *F*_*1*_ = 5.8, *P*<0.001; caste × colony: *F*_*2*_ = 1.4, *P* = 0.08, Fig. S4). We also detected significant differences in the parabasalid community among castes and between colonies (caste: *F*_*2*_ = 5.6, *P*<0.001; colony: *F*_*1*_ = 2.7, *P* = 0.01; caste × colony: *F*_*2*_ = 1.4, *P* = 0.15, Fig. S4), but the difference between colonies was less clear compared with the oxymonad community based on NMDS (Fig. S4). PERMANOVA of nymphs and alates with colony, caste, and sex as explanatory variables showed no significant effect of individual sex on either the oxymonad (*F*_*1*_ = 1.11, *P* = 0.33) or parabasalid (*F*_*1*_ = 1.1, *P* = 0.33) community composition. All oxymonad protists were detected in workers, whereas five species were not detected in several samples of nymphs and alates (Fig. S4, Table S2). Among nymphs, the detection ratios of *D. leidyi, D. porteri* type I, and *D. parva* were 0.947, 0.947, and 0.579, respectively, and among alates, the ratios of *P. modesta, D. rugosa, D. porteri* type I, and *D. parva* were 0.772, 0.944, 0.778, and 0.722, respectively (Table S2). Among parabasalid protists, *Tr. agilis* and *Te. mirabilis* were detected in all nymphs and alates and most workers, whereas the detection rates of *Holomastigotes* sp. or *Microjoenia* sp. were lower in alates and nymphs than in workers. The detection rates of unclassified Trichomonadidae 1 and 2 were relatively low in all castes compared to other parabasalids (Table S3).

Total body weight, body weight without the gut, and gut weight differed significantly among workers, nymphs, A2, A7, and A7w (LMM, LRT, 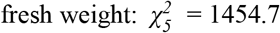, *P*<0.001; 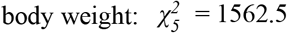, *P*<0.001; 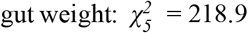, *P*<0.001, Fig. S5). Nymphs and alates showed higher fresh weight and body weight than workers, whereas the gut weight of workers was similar in A2 but heavier than that of nymphs, A7, and A7w (Fig. S5). Among A2 and A7w nymphs, females showed higher fresh weight and body weight than males, whereas the gut weight did not differ significantly between sexes (Fig. S5).

### (b) No transmission of protists from workers to alates

After keeping protist-free alates with workers for 10 days, no protists were observed from any alate gut samples (Fig. 4). Therefore, after eclosion, alates do not obtain protists from workers via proctodeal trophallaxis. In contrast, all post-moult workers regained protists in the hindgut (Fig. 4).

**Fig. 4.**
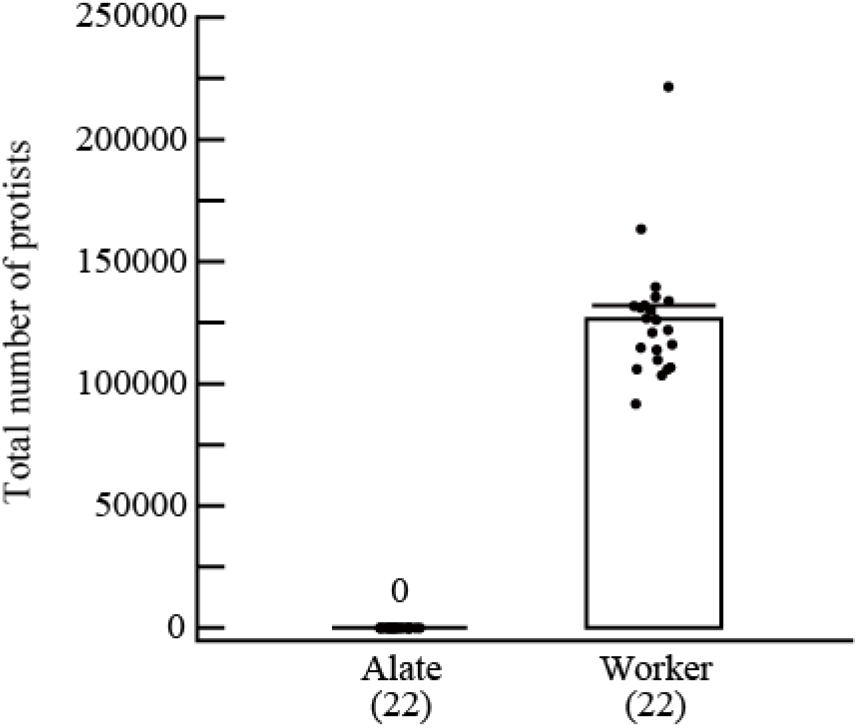
No transmission of intestinal protists from workers to alates after eclosion The total number of protists in alates and workers after moulting. Points show individual measurements. Error bars denote the standard error. The number of samples used is indicated in parentheses.

### (c) Changes in the community composition of nymphs contribute to transmission efficiency

Figure 5a shows the transmission efficiency (i.e., the proportion of hypothetical alates with all protist species) based on two scenarios: transmission of protist communities from workers or nymphs. The minimum number of protists cells required to ensure that all protist species would be transmitted (i.e., transmission efficiency of 100%) differed between the two scenarios: 3500 and 6500 protist cells were required to be transmitted from nymphs and workers, respectively. This difference was attributed to the observation that *Te. mirabilis*, which exhibited the lowest proportion of the protist community, comprised a greater proportion of the community in the gut of nymphs than in that of workers. A comparison with the empirical data (Fig. 5b) showed that most A7 alates (17/20) harboured >3500 protist cells, but very few individuals (3/20) harboured >6500 cells. This result suggests that in the case of random protist transmission from workers, some alates would lack one or more protist species. Therefore, changes in the composition of the protist community in nymphs would be important for increasing the transmission efficiency of the whole protist community.

**Fig. 5.**
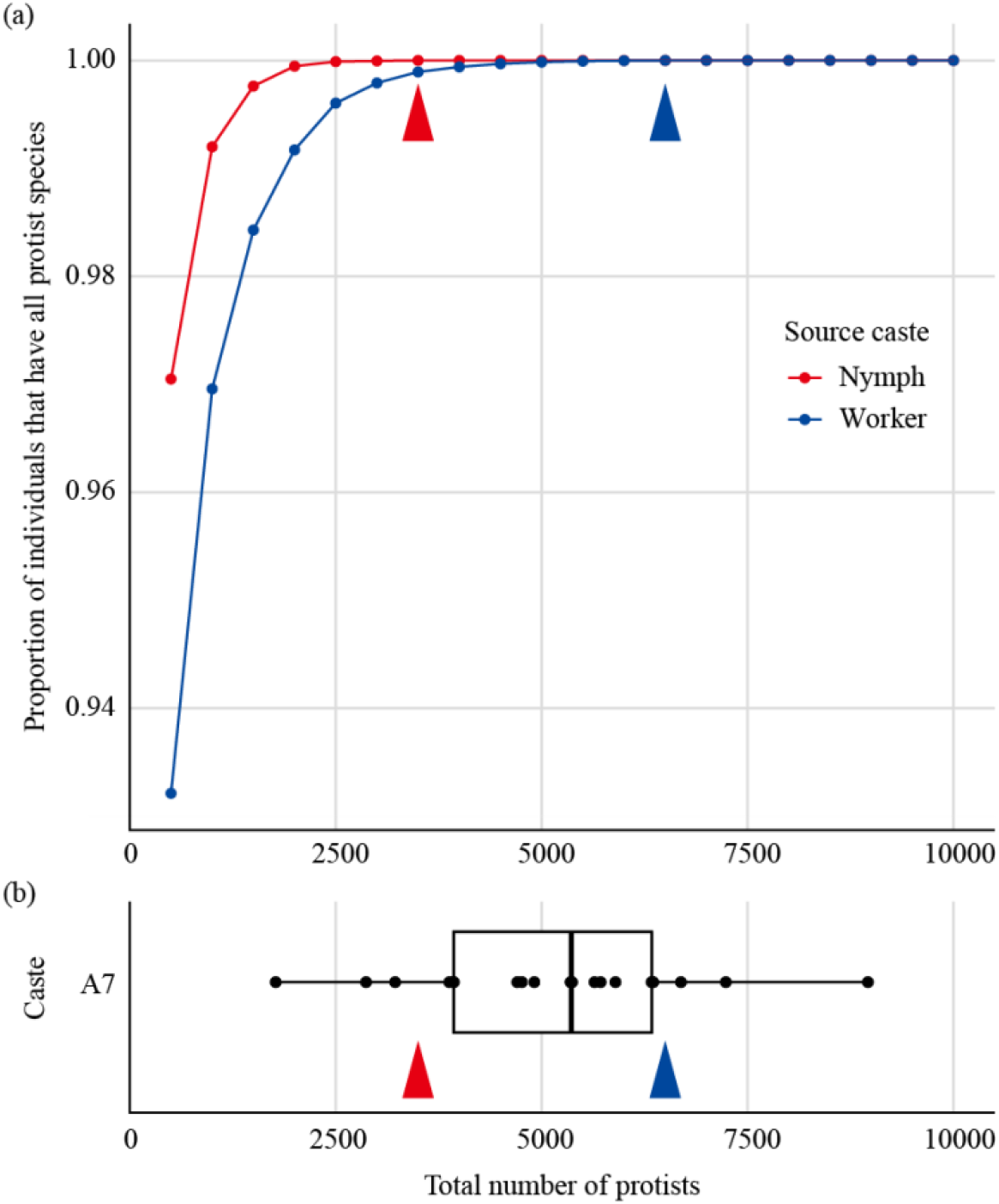
Effect of changes in the protist community composition of nymphs on transmission efficiency (a) Simulation of the number of protists required to transmit all species of protists from nymphs or workers to alates. (b) Boxplot showing the total number of protists in alates (A7: alate isolated 7 days after eclosion). Points are individual measurements, and boxplot shows the range, median, and quartiles for each treatment. Red and blue arrowheads indicate the minimum number of protist cells required to harbour all protist species in the hypothetical alate in two scenarios: random transmission of protists from either nymphs or workers, respectively.

### (d) Protist species–specific changes in cell morphology during alate ecdysis

Our preliminary observation showed that the morphology of *Dinenympha leidyi* changes significantly during adult eclosion, whereas that of *D. exilis* does not. To evaluate the degree of morphological changes in these two species during adult eclosion, we designed species-specific probes, observed each protist species using FISH, and quantified various morphologic parameters (Fig. S6). With the exception of the area of *D. exilis* in colony IV, the area and circularity of both *D. leidyi* and *D. exilis* changed significantly during adult eclosion (Fig. S7). In the gut of workers, the size of *D. leidyi* cells varied widely, and many cells were slightly twisted (Figs. S6, S7). During nymph-adult eclosion, the cells became much smaller and more circular (Figs. S6, S7), suggesting that they were strongly folded during this period. After eclosion, the cells became larger, and their shape returned to a form more like that of workers (Figs. S6, S7). In contrast, few changes were observed for *D. exilis* (Figs. S6, S7). These cells almost always maintained an elongated shape, even though the cells were rounded slightly in the gut of nymphs (Figs. S6, S7). No apparent changes in the morphology of the parabasalid species *Tr. agilis* and *Te. Mirabilis* were observed during termite adult eclosion.

## Discussion

We characterised the caste-specific dynamics of the protist community during development of the termite *R. speratus*. In worker-worker moults, no protists were retained during moulting (Fig. 3a), and the workers obtained protists from nestmates (Fig. 4). In contrast, in nymph-adult eclosion, individual termites retained almost all protists (Fig. 3b, c) and never obtained them from workers (Fig. 4). Therefore, the protist community of nymphs is the primary determinant of the community of alates, and maintenance of the protist community during eclosion is essential for successful transmission. Our results also showed that the number of protists in nymphs and alates was dramatically lower than that of workers (Fig. 3a, b), likely because alates must reduce their weight for further dispersal (Fig. S5).

In nymphs and alates, the proportions of *Tr. agilis* and *Te. mirabilis*, which exhibited the lowest proportions in workers, increased markedly (Figs. 3c, S1). Our simulation-based analysis revealed that such changes in the protist community in nymphs enhance the transmission efficiency of whole protist communities by reducing the risk of losing these protists (Fig. 5). Both of the abovementioned protist species beneficially affect the host: *Tr. agilis* and *Te. mirabilis* cells are large (Fig. 1g, h) and can therefore digest large wood particles via cellulase [56, 57]. In addition, the symbiotic bacteria associated with both protist species contribute to the termite-gut ecosystem by supplementing various nitrogenous compounds and by preventing the accumulation of molecular hydrogen, which otherwise would suppress the fermentation of cellulose [13, 58, 59]. The results of the present study suggest that changes in the composition of the protist community in nymphs and maintenance of the composition until alate dispersal play important roles in ensuring the successful transmission of these protist species.

By contrast, our amplicon sequencing analysis revealed decreases in the ratios of several protist species other than *Tr. agilis* and *Te. mirabilis* in nymphs and alates compared to workers (Tables S2, S3), suggesting that not all protist species are completely maintained throughout the transmission process. This result was consistent with previous studies examining *Reticulitermes grassei* [23] and *Coptotermes* sp. [25]. Velenovsky et al. (2023) suggested that biparental transmission contributes significantly to the high rates of occurrence of protist species in field colonies of *Coptotermes* sp. In termites, alates usually form monogamous pairs to establish a new colony, and both founders engage in foraging, reproduction, and caring for larvae via anus-to-mouth feeding [60]. Previous research showed that the number of protists in both male and female founders increases dramatically after colony founding [61-63]. As no significant sex-related differences in the number or community composition of protists or the weight of the gut in nymphs and alates were observed in the present study, the contribution of both sexes as carriers of the protist community would be essentially equal. In addition to the increase in rare protist species before dispersal that we showed in the present study, mixing protist communities from male and female alates during colony founding may improve the transmission efficiency of all protist species in the community.

Our results showed that maintenance of protist communities during eclosion is essential for successful transmission. As the conditions in the hindgut can change dramatically during adult eclosion due to shedding of the epithelium of the hindgut wall [27], protists may have to deal with unusual environmental changes during eclosion. We observed synchronised morphological changes in certain protist species during adult eclosion of termites. The size of *D. leidyi* cells decreased, and the cells appeared to be strongly folded just after moulting (Figs. S6, S7). The cells returned to their normal shape in the alates by 7 days after eclosion (Figs. S6, S7). Such changes during host eclosion have been reported in various termite species [20, 29, 64]. For instance, cells of the oxymonad protist *Streblomastix strix* harboured by termites of the genus *Zootermopsis* usually exhibit an elongated morphology, but they exhibit a rounded shape during adult eclosion [29]. These observations suggest that morphological changes in some protist species enable them to withstand the environmental changes in the hindgut associated with adult eclosion. In contrast, the morphology of *D. exilis* exhibited only slight changes during adult eclosion (Figs. S6, S7). Strategies for overcoming environmental changes may differ between protist species. Although why and how such species-specific morphological changes are induced during adult eclosion remain to be investigated, future studies focusing on physiological changes in the gut environment and protists might address these questions.

Protist communities in the termite gut are maintained within colonies and between generations. Our empirical data and simulation revealed that increasing the proportions of species that are few in number increases their transmission efficiency during dispersal. Such increases in rare species would occur every generation, preventing their disappearance due to significant bottlenecks. This might help maintain the metabolic function of gut microbiota in termites, likely because the role of each protist differs, as shown in the termite *Coptotermes formosanus* [65]. Further studies examining the relationships among protists in the termite gut would enhance our understanding of the mechanisms underlying the maintenance of protist communities in termite life history.

## Supporting information

Electronic Supplemental Materials

## Data accessibility

Data for protist cell counts and R code for simulation are available at https://github.com/TatsuyaInagaki2/Data_alate_protist_community. Additional information is provided in the electronic supplementary material. The demultiplexed MiSeq data sets have been deposited in the DDBJ Sequence Read Archive under the BioProject PRJDB15577 with BioSample accession numbers SAMD00589838, SAMD00589839, SAMD00589840, SAMD00589841, SAMD00589842, SAMD00598142, SAMD00598143, SAMD00598144, SAMD00598145, SAMD00598146, SAMD00598147, SAMD00598148, SAMD00598149, SAMD00598150, SAMD00598151, SAMD00598152, SAMD00598153, SAMD00598154, SAMD00598155, SAMD00598156, SAMD00598157, SAMD00598158, SAMD00598159, SAMD00598160, SAMD00598161, SAMD00598162, SAMD00598163, SAMD00598164, SAMD00598165, SAMD00598166, SAMD00598167, SAMD00598168, SAMD00598169, SAMD00598170, SAMD00598171, SAMD00598172, SAMD00598173, SAMD00598174, SAMD00598175, SAMD00598176, SAMD00598177, SAMD00598178, SAMD00598179, SAMD00598180, SAMD00598181, SAMD00598182, SAMD00598183, SAMD00598184, SAMD00598185, SAMD00598186.

## Authors’ contributions

T.I.: conceptualization, data curation, formal analysis, funding acquisition, investigation, methodology, validation, visualization, writing—original draft; K.I.: methodology, writing—review and editing; K.T.: investigation, methodology, writing—review and editing; Y.H.: conceptualization, funding acquisition, supervision, writing—review and editing.

## Acknowledgments

We thank Daiki Kato, Mamoru Takata and members of Hongoh Lab at Tokyo Institute of Technology for the helpful discussion. We also would like to thank FORTE SCIENCE COMMUNICATIONS for English language editing.

