## Supplementary material for "Transmission dynamics of symbiotic protist communities in the termite gut: association with host adult eclosion and dispersal": Electronic Supplemental Materials

**The file includes:**

- Table S1 Protist community composition in hypothetical workers and nymphs for the simulation of vertical transmission
- Fig. S1 Proportions of *Trichonympha agilis* and *Teranympha mirabilis*
- Fig. S2 Phylogenetic tree of oxymonad protists and 20 OTUs derived from amplicon sequencing
- Fig. S3 Phylogenetic tree of parabasalid protists and OTUs derived from amplicon sequencing
- Fig. S4 Protist community of each caste determined by amplicon sequencing
- Table S2 Detection rate of oxymonad protists from each caste based on amplicon sequencing
- Table S3 Detection rate of parabasalid protists from each caste based on amplicon sequencing
- Fig. S5 Gut and body weights of workers, nymphs, and alates
- Fig. S6 Specific detection of two oxymonad protist species by fluorescence in situ hybridization in each caste
- Fig. S7 Morphological changes in two oxymonad protist species during alate eclosion

Table S1 Protist community composition in hypothetical workers and nymphs for the simulation of vertical transmission. The numbers were determined as the average of empirical data.

| Caste | Protist groups | | | | |
|---|---|---|---|---|---|
| | *Pyrsonympha* spp. | *Dinenympha exilis* | *D. porteri* type III | *D.* spp.1 | *D.* spp.2 |
| Nymph | 819 | 1914 | 1310 | 2060 | 1509 |
| Worker | 10415 | 18618 | 6009 | 20033 | 15815 |

| Caste | Protist groups | | | |
|---|---|---|---|---|
| | *Holomastigotes elongatum* | *Trichonympha agilis* | *Teranympha mirabilis* | Small Protists |
| Nymph | 185 | 210 | 24 | 959 |
| Worker | 1949 | 374 | 107 | 5924 |

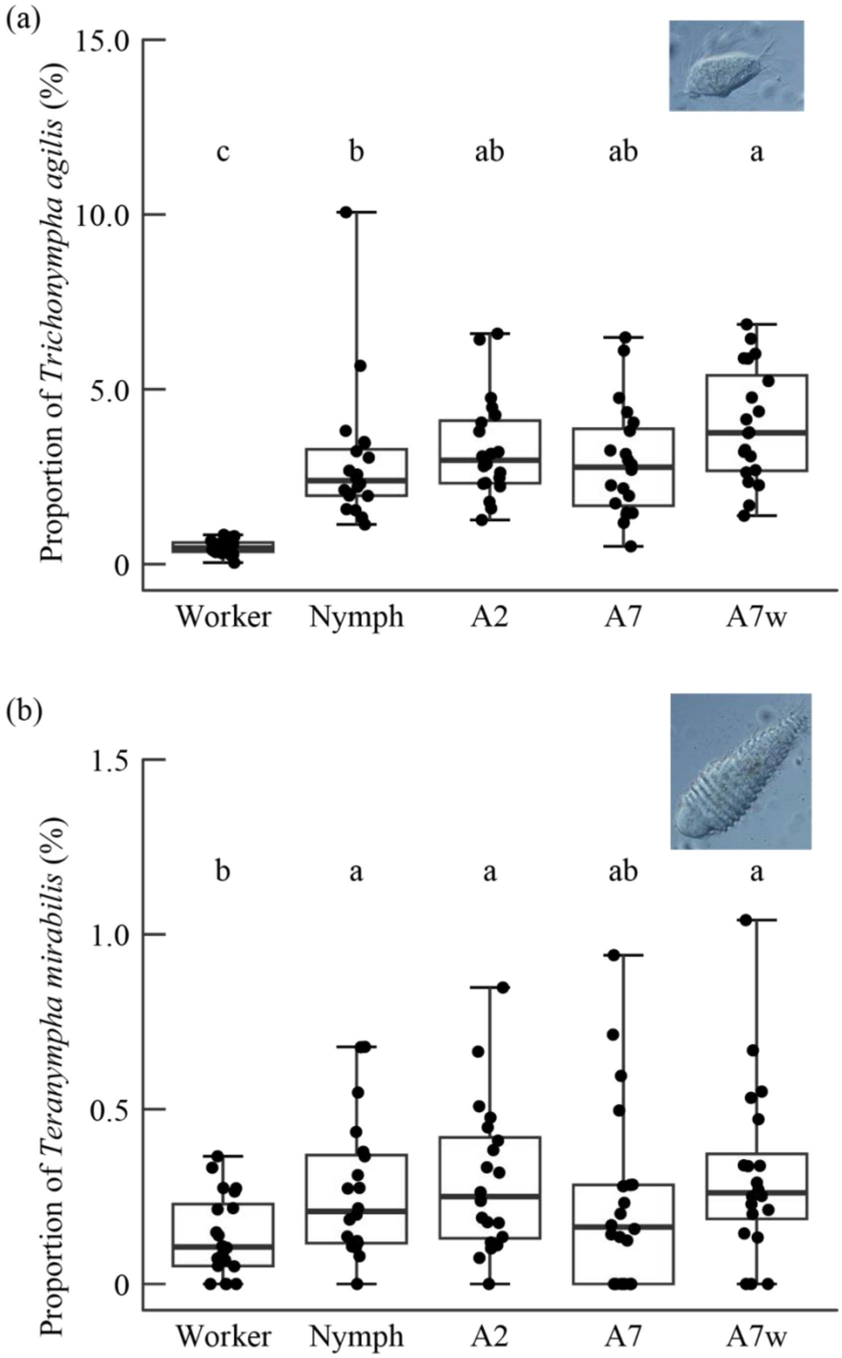


Fig. S1 Proportions of *Trichonympha agilis* and *Teranympha mirabilis*

Proportions of two protist species in each caste/treatment derived from cell counts, indicated by percentage. Points are individual measurements, and boxplots show the range, median, and quartiles for each treatment. Different characters denote significant differences (Tukey's HSD test after GLM with quasi-binomial error distribution, $P<0.05$).

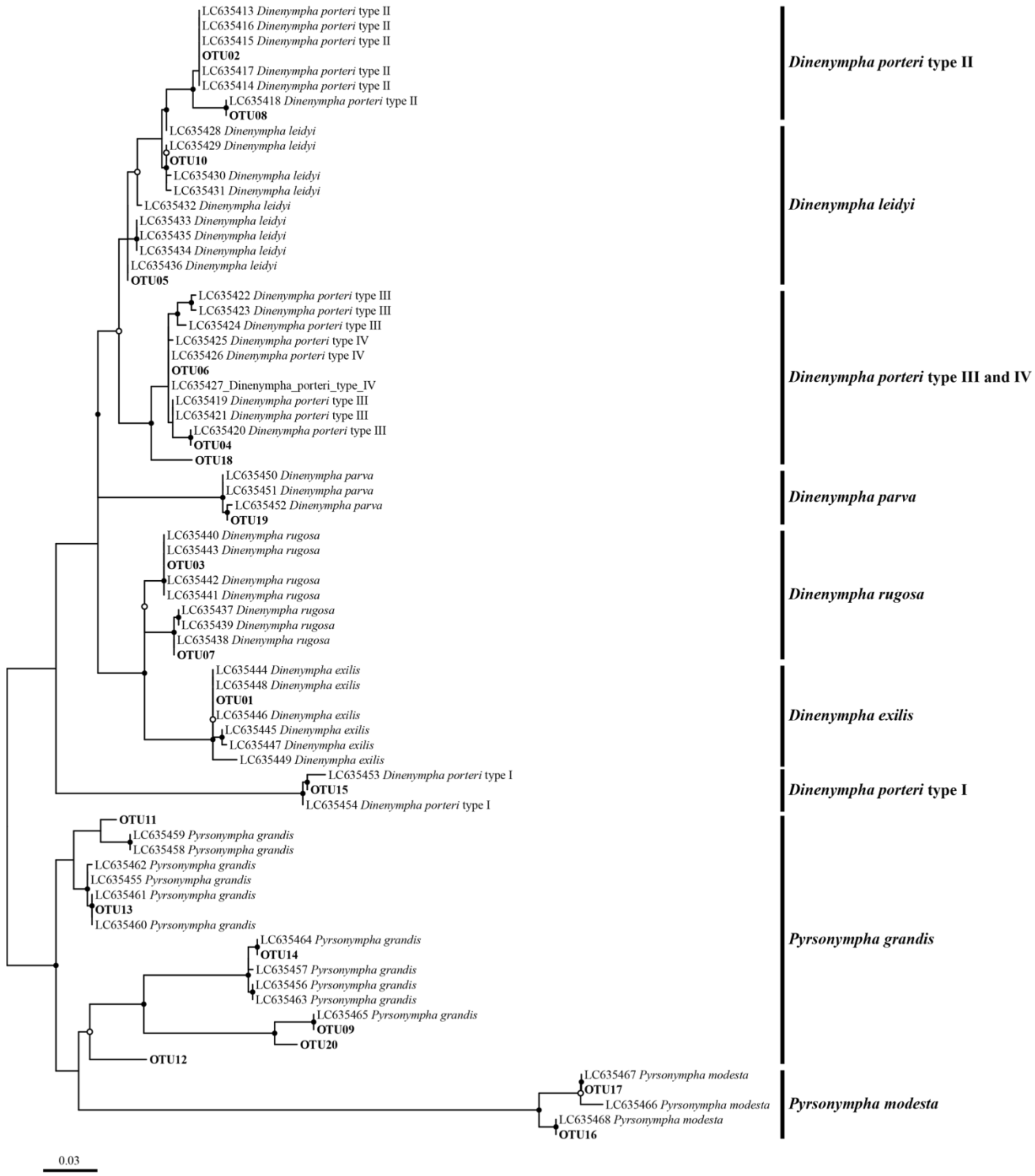


Fig. S2 Phylogenetic tree of oxymonad protists and 20 OTUs derived from amplicon sequencing

The tree was constructed based on 417 aligned nucleotide sites of the V3–V4 region of 18S rRNA genes. Sequences of 20 OTUs in this study and 56 single oxymonad cells from a previous study [1] were used. Assigned species of oxymonads are indicated with solid lines. Branch support is indicated as filled circles (ultrafast bootstrap ≥90%) or open circles (ultrafast bootstrap ≥80%).

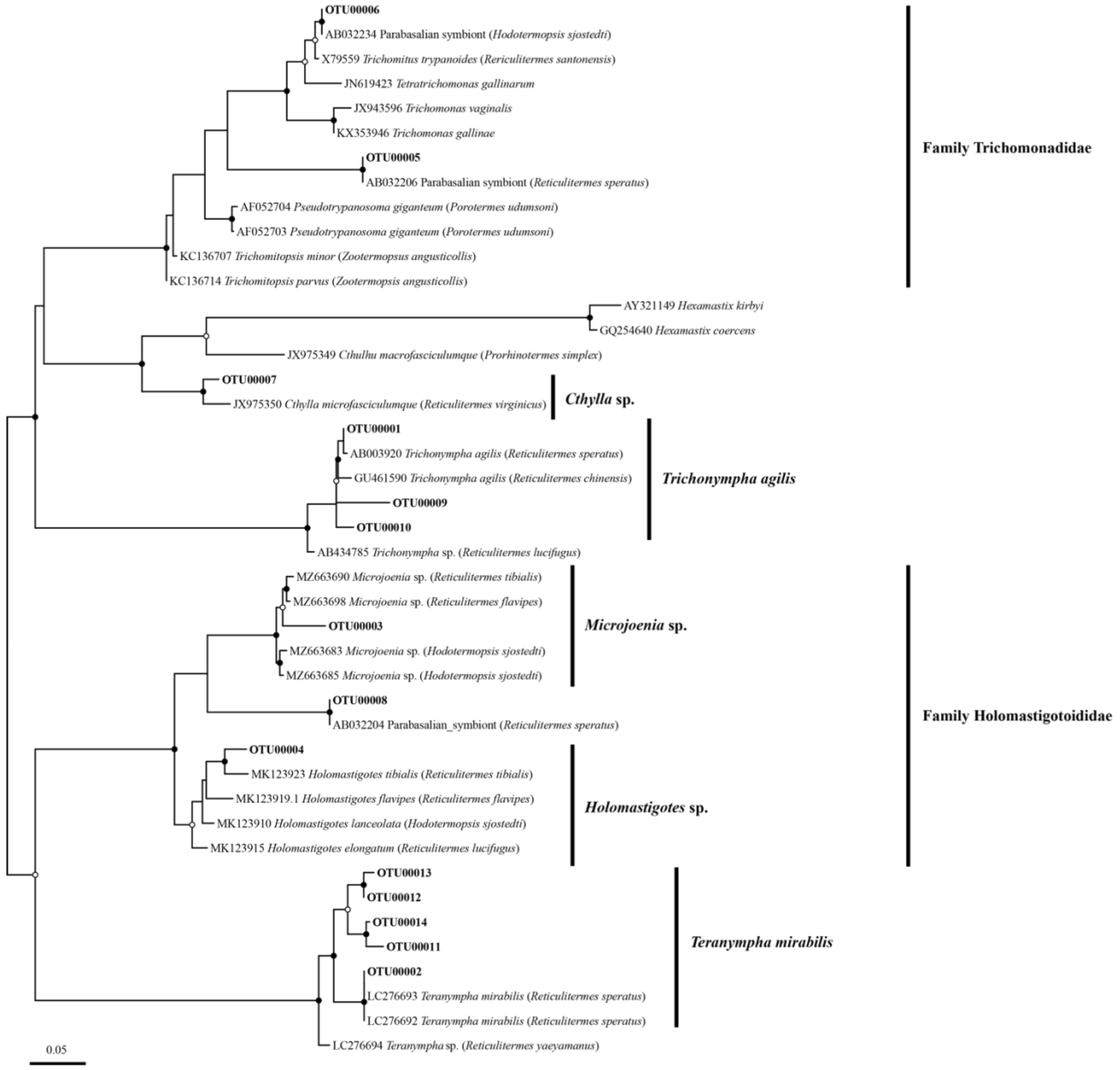


Fig. S3 Phylogenetic tree of parabasalid protists and OTUs derived from amplicon sequencing

The tree was constructed based on 352 aligned nucleotide sites of the V3–V4 region of 18S rRNA genes. Sequences of 10 OTUs in this study and other parabasalids from public databases were used. Each OTU was assigned to species-level taxa as follows: *Trichonympha agilis* (OTU 01, 09), *Teranympha mirabilis* (OTU 02, 10), *Holomastigotes* sp. (OTU 04), *Microjoenia* sp. (OTU 03), unclassified Holomastigotoididae (OTU 08), unclassified Trichomonadidae 1 (OTU 05), unclassified Trichomonadidae 2 (OTU 06), or *Cthylla* sp. (OTU 07). Branch support is indicated as filled circles (ultrafast bootstrap ≥90%) or open circles (ultrafast bootstrap ≥80%).

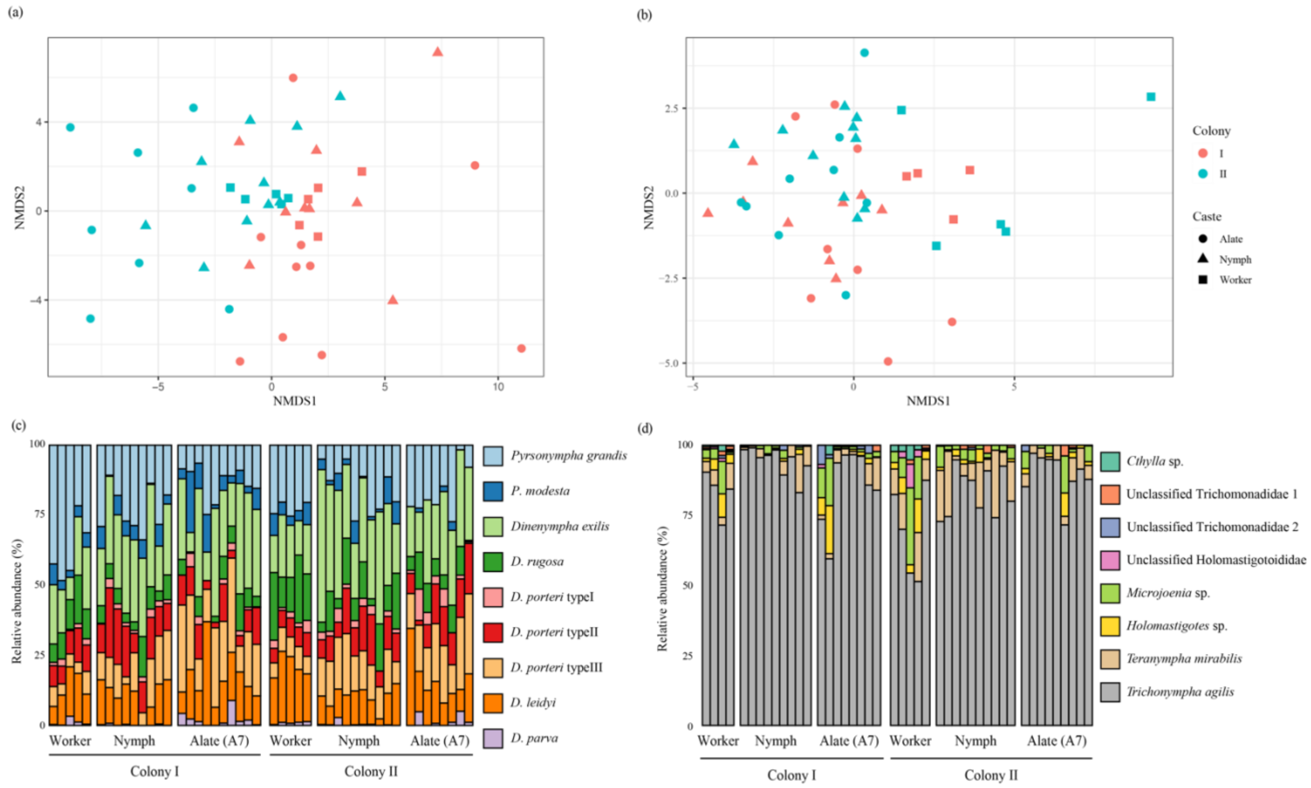


Fig. S4 Protist community of each caste determined by amplicon sequencing

Upper graphs show NMDS plots based on the Euclidean distance between samples calculated from centred log-ratio transformed data, and bottom graphs show bar plots for (a, c) oxymonad and (b, d) parabasalid protist communities.

Table S2 Detection rate of oxymonad protists from each caste based on amplicon sequencing

| Caste | N | Oxymonad protists | | | | |
|---|---|---|---|---|---|---|
| | | *Pyrsonympha grandis* | *P. modesta* | *D. exilis* | *D. rugosa* | *D. leidyi* |
| Worker | 10 | 1.000 | 1.000 | 1.000 | 1.000 | 1.000 |
| Nymph | 19 | 1.000 | 1.000 | 1.000 | 1.000 | 0.947 |
| Alate (A7) | 18 | 1.000 | 0.722 | 1.000 | 0.944 | 1.000 |

| Caste | N | Oxymonad protists | | | |
|---|---|---|---|---|---|
| | | *D. porteri* type I | *D. porteri* type II | *D. porteri* type III | *D. parva* |
| Worker | 10 | 1.000 | 1.000 | 1.000 | 1.000 |
| Nymph | 19 | 0.947 | 1.000 | 1.000 | 0.579 |
| Alate (A7) | 18 | 0.778 | 1.000 | 1.000 | 0.722 |

Table S3 Detection rate of parabasalid protists from each caste based on amplicon sequencing

| Caste | N | Parabasalid protists | | | |
|---|---|---|---|---|---|
| | | *Trichonympha agilis* | *Teranympha mirabilis* | *Holomastigotes* sp. | *Microjoenia* sp. |
| Worker | 9 | 1.000 | 0.889 | 1.000 | 1.000 |
| Nymph | 19 | 1.000 | 1.000 | 0.789 | 0.842 |
| Alate(A7) | 17 | 1.000 | 1.000 | 0.765 | 0.824 |

| Caste | N | Parabasalid protists | | | |
|---|---|---|---|---|---|
| | | Unclassified Holomastigotoididae | *Cthylla* sp. | Unclassified Trichomonadidae 1 | Unclassified Trichomonadidae 2 |
| Worker | 9 | 0.889 | 0.889 | 0.556 | 0.333 |
| Nymph | 19 | 0 | 0.105 | 0.368 | 0.211 |
| Alate(A7) | 17 | 0.059 | 0.059 | 0.412 | 0.412 |

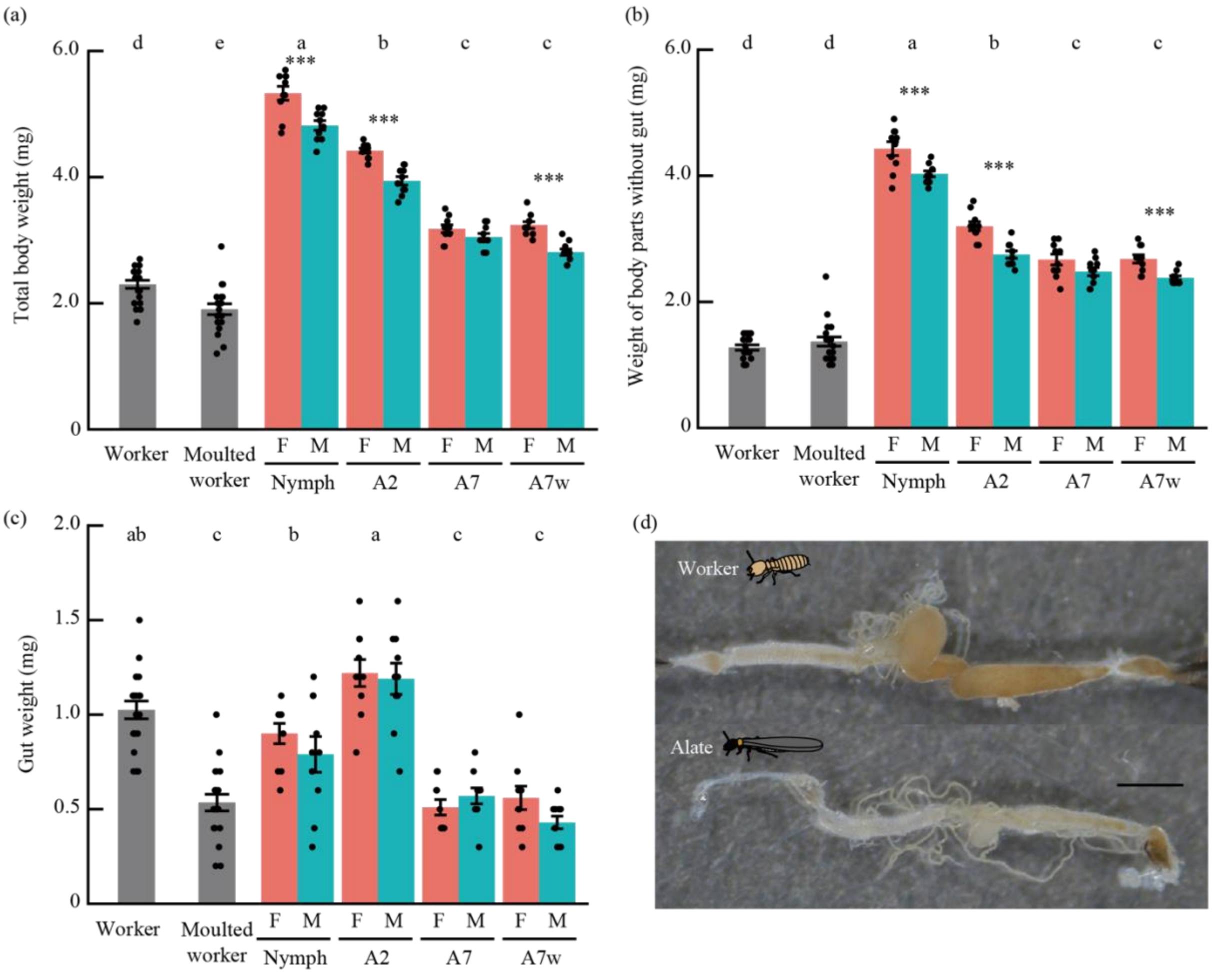


Fig. S5 Gut and body weights of workers, nymphs, and alates

(a) Fresh weight of the whole body, including the gut, (b) body weight except for the gut, and (c) gut weight of workers, nymphs, and alates. Error bars denote the standard error, and points show individual measurements. Different characters denote significant differences (Tukey's HSD after LMM, colony was included as a random factor, $P<0.05$). Asterisks show significant differences between sexes in nymphs and alates (LMM, colony was included as a random factor, Bonferroni correction, corrected $\alpha = 0.0125$). (d) Whole gut of a worker (upper) and an alate (bottom). Scale bar: 1 mm.

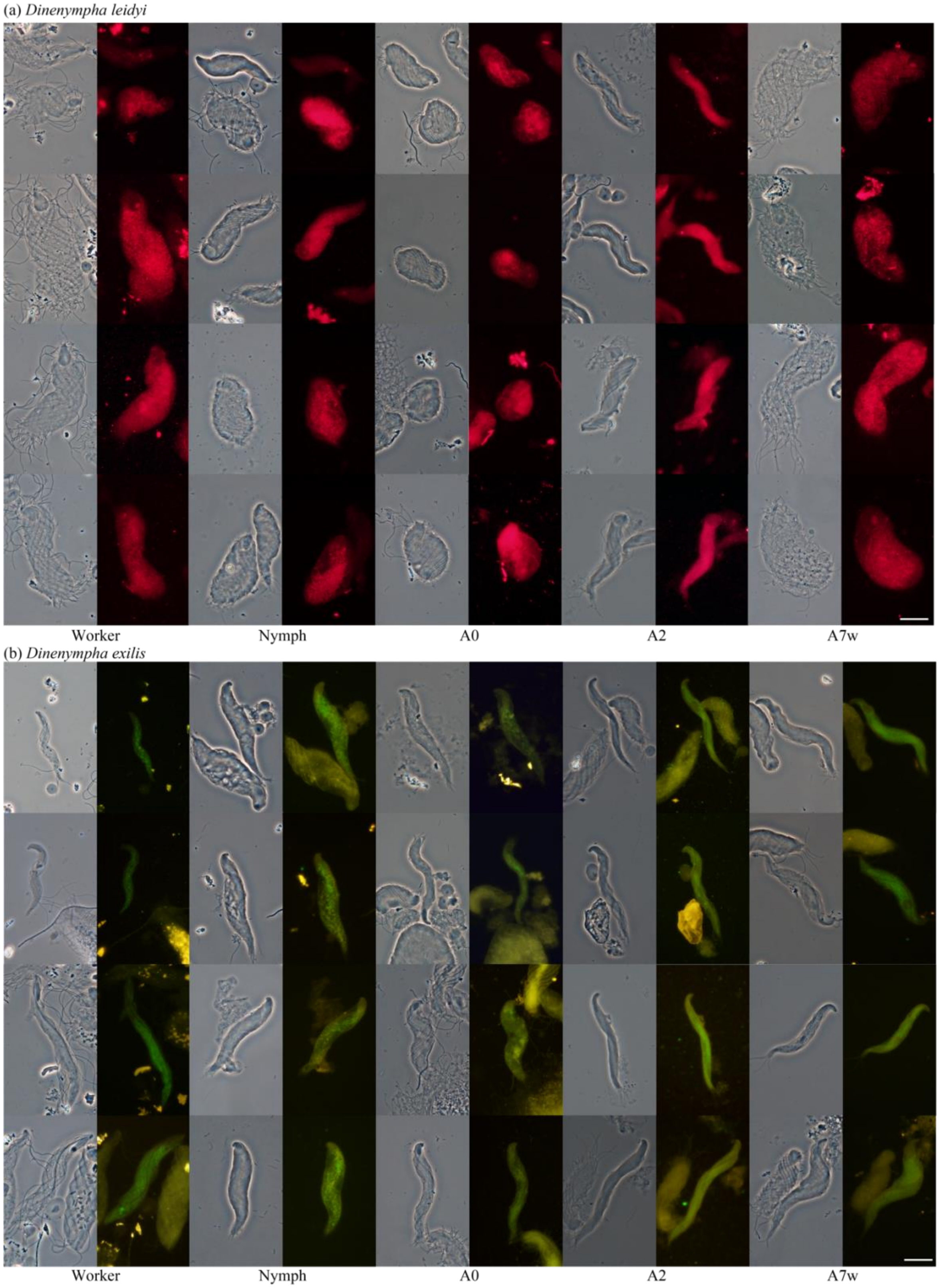


Fig. S6 Specific detection of two oxymonad protist species by fluorescence in situ hybridization in each caste Phase-contrast images and specific detection of (a) *Dinenympha leidyi* with a TEX-labelled probe (red) and (b) *D. exilis* with a FAM-labelled probe (green). Left to right: worker, nymph, A0 (alates just after eclosion), A2 (alates isolated 2 days after eclosion), and A7w (alates approximately 7 days after eclosion that were kept with workers). Scale bars in (a) and (b): 20 µm.

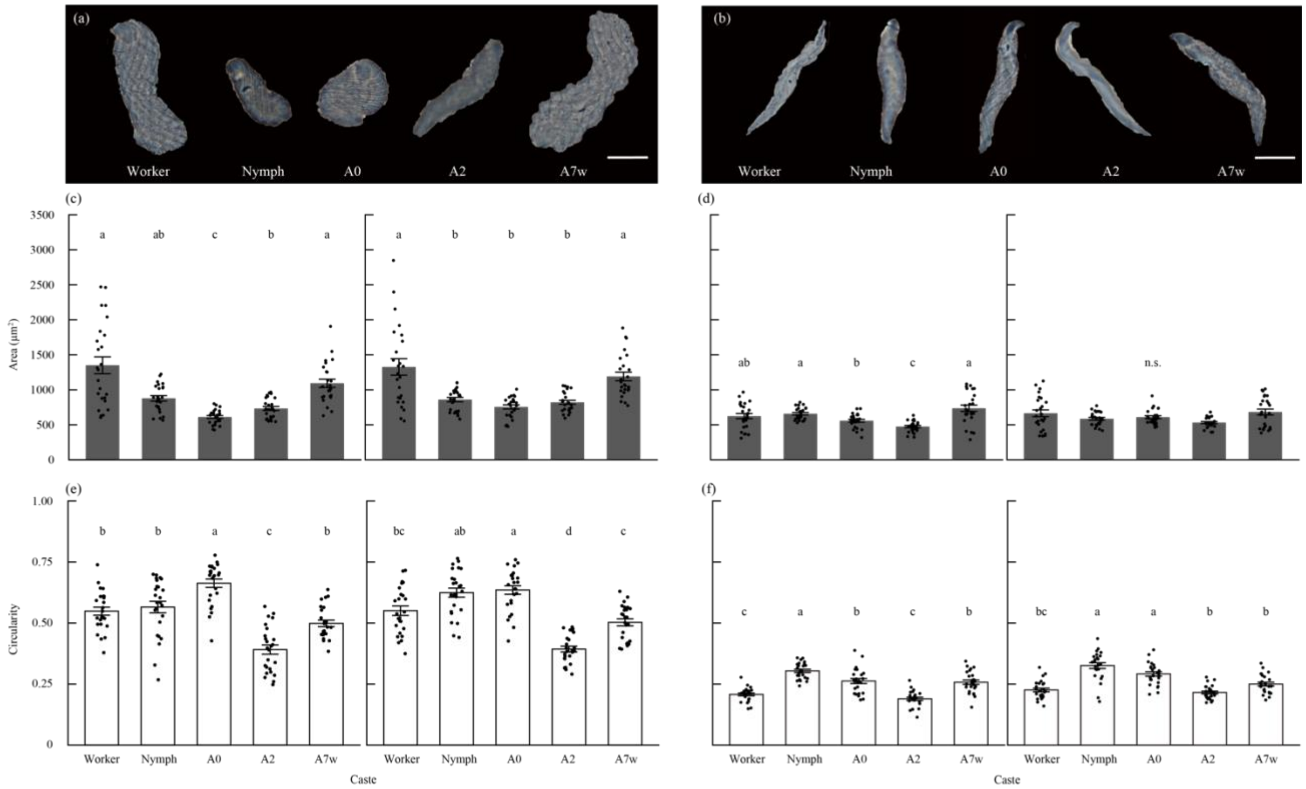


Fig. S7 Morphological changes in two oxymonad protist species during alate eclosion

Images of cells of (a) *Dinenympha leidyi* and *D. exilis* (b) trimmed using the Quick Selection Tool in Adobe Photoshop CC. Scale bar: 20 µm. The area (c, d) and circularity (e, f) of each protist species in each caste from two colonies are shown. In each panel, left and right graphs show colonies III and IV, respectively. Error bars denote the standard error. Different characters indicate significant differences in each colony (pairwise Wilcoxon rank-sum test, Bonferroni correction, corrected $\alpha = 0.005$).
